# Multiplex imaging combined to machine learning enable automated profiling of cortical malformations: applications in tuberous sclerosis complex

**DOI:** 10.1101/2025.11.24.690101

**Authors:** Reyes Castaño-Martín, Alice Metais, Francesco Carbone, Naziha Bakouh, Estelle Balducci, Damien Conrozier, Sofian Ameur, Pascale Varlet, Rima Nabbout, Edor Kabashi, Mark Zaidi, Thomas Blauwblomme, Sorana Ciura

**Affiliations:** Translational Research in Neuroscience Lab, Institut Imagine, Université Paris Cité, INSERM U1163, 75015 Paris, France; Institute of Psychiatry and Neuroscience of Paris (IPNP), Université Paris Cité, INSERM U1266, 75014 Paris, France; Service de Neuropathologie, GHU-Paris Psychiatrie et Neurosciences, Hôpital Sainte Anne, F-75014 Paris, France; Bioinformatic platform, Institut Imagine, Inserm UMR1163, 75015 Paris, France; Department of Biological hematology, Assistance Publique Hôpitaux de Paris, 75015 Paris, France; Histology platform, SFR Necker, INSERM US24, CNRS UAR3633, 75015 Paris, France; Université de Paris Cité, Paris, France; Department of Pediatric Neurology, Hôpital Necker, Assistance Publique Hôpitaux de Paris, 75015 Paris, France; Department of Pediatric Neurosurgery Hôpital Necker, Assistance Publique Hôpitaux de Paris, 75015 Paris, France

**Keywords:** malformation of cortical development, multiplex imaging, machine learning, spatial analysis

## Abstract

Malformations of cortical development such as tuberous sclerosis complex arise within a heterogeneous cellular landscape that conventional histopathology only partially resolves. Here, we combined a 19-marker multiplex immunofluorescence panel with a machine learning-driven image analysis pipeline to map and quantify over 365 000 cells from paediatric surgical cortex, defining the single-cell architecture of TSC lesions. Microtubers were objectively delineated by vimentin and detected in all TSC samples but absent from non-dysplastic controls. Within these structures, balloon cells exhibiting strong pS6 activation occupied lesion cores and were confined to microtuber boundaries, whereas dysmorphic neurons were more diffusely distributed into adjacent cortex. The microtuber niche was dominated by astroglial remodeling: immature and reactive vimentin-positive astrocytes, including Lamp5-positive subsets, accumulated at and around lesion rims, while mature GFAP-positive astrocytes showed only modest changes. Distance-based spatial analyses revealed neuronal exclusion from microtuber centres with gradual recovery in surrounding tissue, indicating local network disruption. Unsupervised clustering and niche modelling recapitulated these spatial gradients, identifying a glial-dominated ecosystem that concentrates balloon cells, increases inter-neuronal distances, and reduces cell-cell interactions. Together, these data support a model in which cortical tubers arise through the coalescence of microtubers orchestrated by balloon cells and reactive gliosis during corticogenesis. Beyond elucidating disease architecture, our automated framework enables reproducible lesion detection, quantitative cell-state mapping, and spatial readouts applicable across malformations of cortical development.

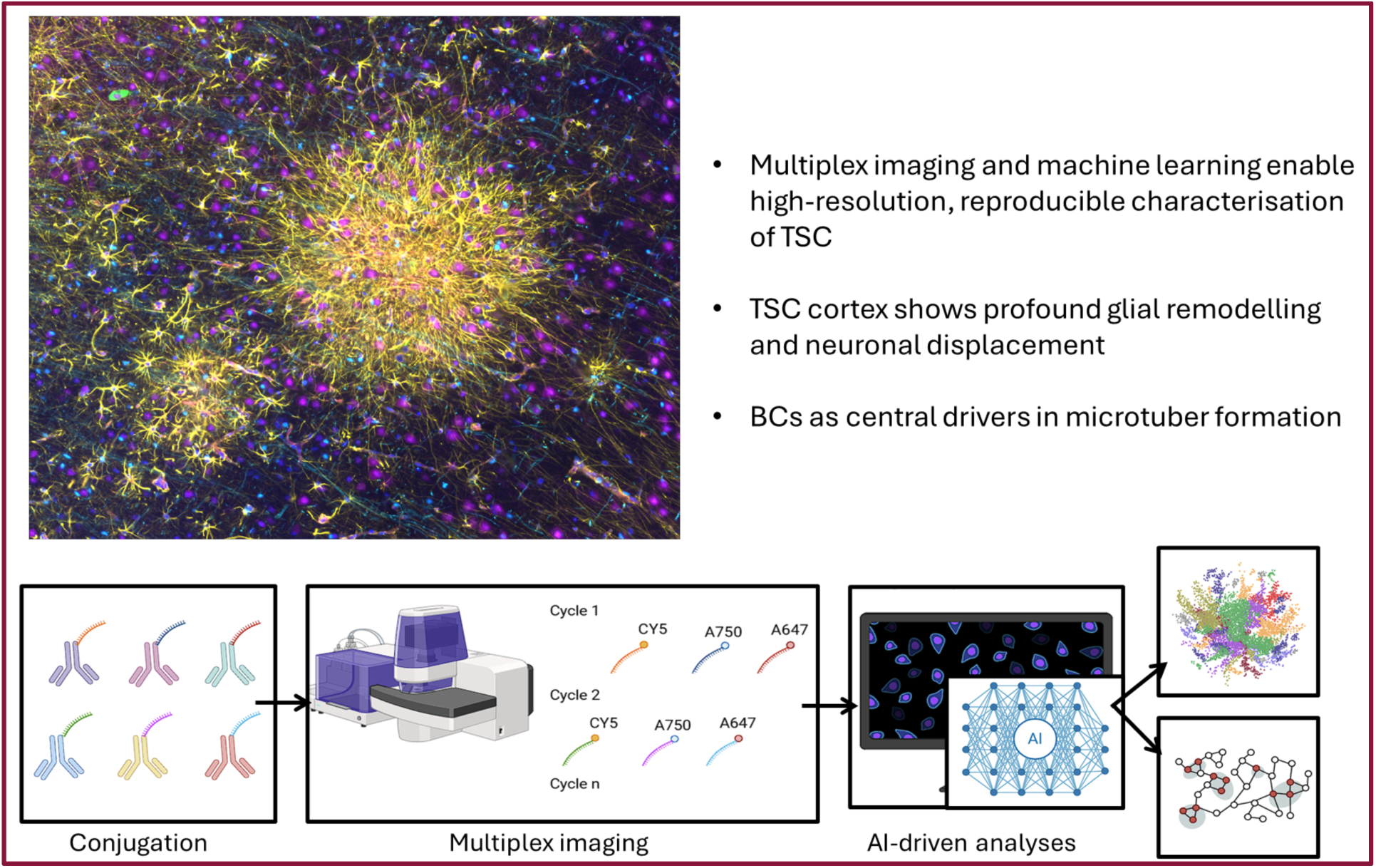

## Introduction

Multiplex spatial imaging and machine learning have revolutionised histopathological analysis by enabling high-resolution, quantitative characterisation of complex tissue architectures ^1–3^. Traditional colorimetric immunohistopathological techniques rely on one or two markers per slide, requiring several tissue sections with possible loss of information when the structure to be observed is exhausted, which limits the ability to capture the intricate cellular interactions and heterogeneity within pathological structures. In contrast, multiplex imaging techniques, such as immunofluorescence or mass cytometry-based approaches, allow for the simultaneous detection of multiple biomarkers within the same tissue section. These advances provide unprecedented spatial context, revealing the cellular composition, molecular expression patterns, and tissue microenvironment ^4^.

However, the vast amount of high-dimensional data generated by multiplex imaging necessitates advanced computational approaches for effective analysis. Machine learning, particularly in the form of feature-based classification and deep learning segmentation models, has become an essential tool for extracting meaningful insights from these datasets. Algorithms trained on annotated histological images can automate cell segmentation, identify pathological features, and classify disease-associated patterns with high accuracy and reproducibility ^5–7^.

Cortical tubers are rare, yet archetypal, malformations of cortical development (MCD), belonging to the mTORopathy group of neuropathologies associated with epilepsy and developmental impairment, that also include Focal Cortical Dysplasia (FCD) type II and Hemimegalencephaly (HME) ^8,9^. Tubers are specifically encountered in patients with Tuberous Sclerosis Complex (TSC) disease, an autosomal dominant disorder associated with mutations in either TSC1 or TSC2 genes affecting 2 million individuals worldwide ^10^. Mutations in TSC genes result in the inactivation of their protein products and subsequent hyperactivation of the mammalian target of rapamycin (mTOR) pathway ^11^. The severity of the disease correlates strongly with the “tuber burden,” defined by the number and size of these tubers in the brain ^12^.

TSC tubers typically range in size from a few millimetres to several centimetres in diameter, and are readily identified by MRI due to their distinct signal and architecture ^13^. The histopathological features of tubers in TSC include abnormally enlarged cells, gliosis, cortical delamination and the loss of specific neuronal subtypes ^14–17^. The enlarged cell types are classified as dysmorphic neurons (DNs) and giant or balloon cells (BCs), with both types displaying strong mTOR hyperactivity ^14^. Tubers are often surrounded by rosette-shaped, satellite-like structures termed microtubers or micronodules that have been less well characterised ^18^. Microtubers are developmental abnormalities found in the periphery of the tubers and thought to be the precursors of the cortical tubers, providing a tool for understanding the very early processes involved in brain malformations in TSC ^16^.

By integrating multiplex imaging combined with machine learning on a cohort of post-surgical biopsies, our work provides for the first time a quantitative, high-resolution spatial analysis of TSC brain biopsies, using a custom antibody-based multiplex imaging and machine learning-driven spatial mapping to systematically characterise the cellular composition and spatial organisation of microtubers. Specifically, we reveal the central role of pS6-positive balloon cells in shaping microtuber architecture, providing evidence that supports a central role of these cells in organising a vimentin-positive glial population that forms a distinctive pathological network disrupting normal cortical organisation.

Distance-based analysis is a crucial tool for understanding cellular interactions within pathological structures, as it allows for the quantification of spatial gradients and heterogeneity in tissue organisation. Previous studies on kidney injuries or cancer have demonstrated its importance in elucidating microenvironmental factors that contribute to disease progression ^6,19^. Using clustering and distance-based analyses, we explore how neuronal and glial interactions within microtubers differ from surrounding cortical tissue, providing novel insights into the spatial determinants of TSC pathology.

All in all, our findings underscore the importance of advanced imaging techniques in unravelling the intricate cellular landscapes and spatial interactions that could be generalised in other cerebral cortex diseases.

## Results

### Clinical data

We chose to develop the first multiplex analysis of MCD cortical tissue on a disease with reproducible and characteristic histological findings to decrease interpatient heterogeneity. Therefore, we focused on epileptic children affected by tuberous sclerosis operated of a cortical tuber. 9 cortical samples from 7 epileptic children were included in this study (Supplementary Table 1): 6 cortical tubers, and 3 control neocortices from the temporal pole adjacent yet remote to hippocampal sclerosis (n=2) or dysembryoplastic neuroepithelial tumour (DNET). Included in the cohort there were 5 girls and 2 boys, mean age at surgery was 5.1 years old (range 1.1-15.1); with a mean follow up of 25 months (range 7-41 months) and 6/7 patients are seizure free. The 5 cortical tubers were encountered in children with sporadic TSC (germline mutation in TSC1 in 2 cases, TSC2 in 3 cases).

### Multiplex neurobiology panel in TSC

We designed and conjugated a novel panel of 19 markers (Supplementary Table 2) to capture the cellular diversity of TSC, obtaining the first concomitant image of multiple markers in a TSC cortical brain tissue (Fig. 1). We first selected markers that allowed us to distinguish between the different cortical cell types (Fig. 2a, b): inhibitory (Gad1) and excitatory (TBR1) neurons (NeuN and MAP2), astrocytes (GFAP), microglia (Iba1 and CD68), oligodendrocytes (Olig2 and MOG) and blood vessels (CD31 and VIP). Based on previous publications ^15^, we decided to decipher the interneuron subtypes such as Calbindin-D28k (CB), Calretinin (CR), Parvalbumin (PV), Somatostatin (SST), Neuropeptide Y (NPY) and Lysosomal associated membrane protein 5 (Lamp5). To identify mTOR activation and tuberous structures we used phospho-S6 ribosomal protein (pS6) and vimentin (Vim), respectively ^20^ (Fig. 2a, b). All tissue and cellular annotation were performed under the close supervision of a neuropathologist (MA). Two brain areas were distinguished, white and grey matter. Due to the high autofluorescence of the white matter, only the grey matter was retained for the downstream analysis. The grey matter was positive for MAP2 and Gad1, while the white matter was distinguished by a strong signal for Olig2 and GFAP (Fig. 1). In the TSC tissue, tuberous structures that included subpial gliosis were distinguished by the presence of glial markers such as GFAP, a visual correlation between vimentin, pS6 and partly Lamp5 was noted, but not with microglial or oligodendrocyte markers such as Iba1 or Olig2 (Fig. 1).

**Fig. 1.**
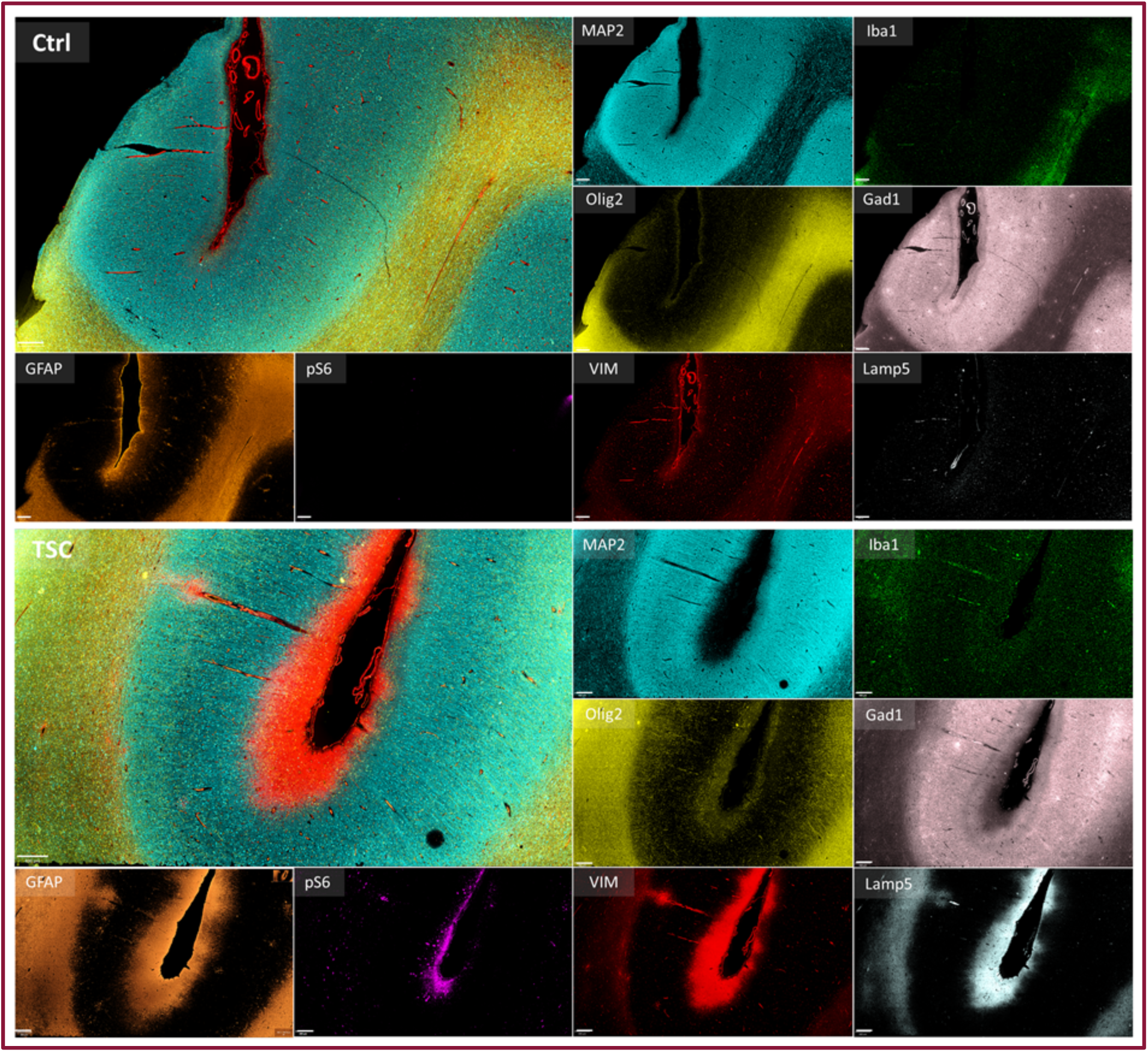
Multiplex image of a comparison of the cortical structure of control and TSC tissue. 3 areas are easily differentiated, grey matter (blue), white matter (yellow) and subpial gliosis (red). Scale bar = 400 µm.

**Fig. 2.**
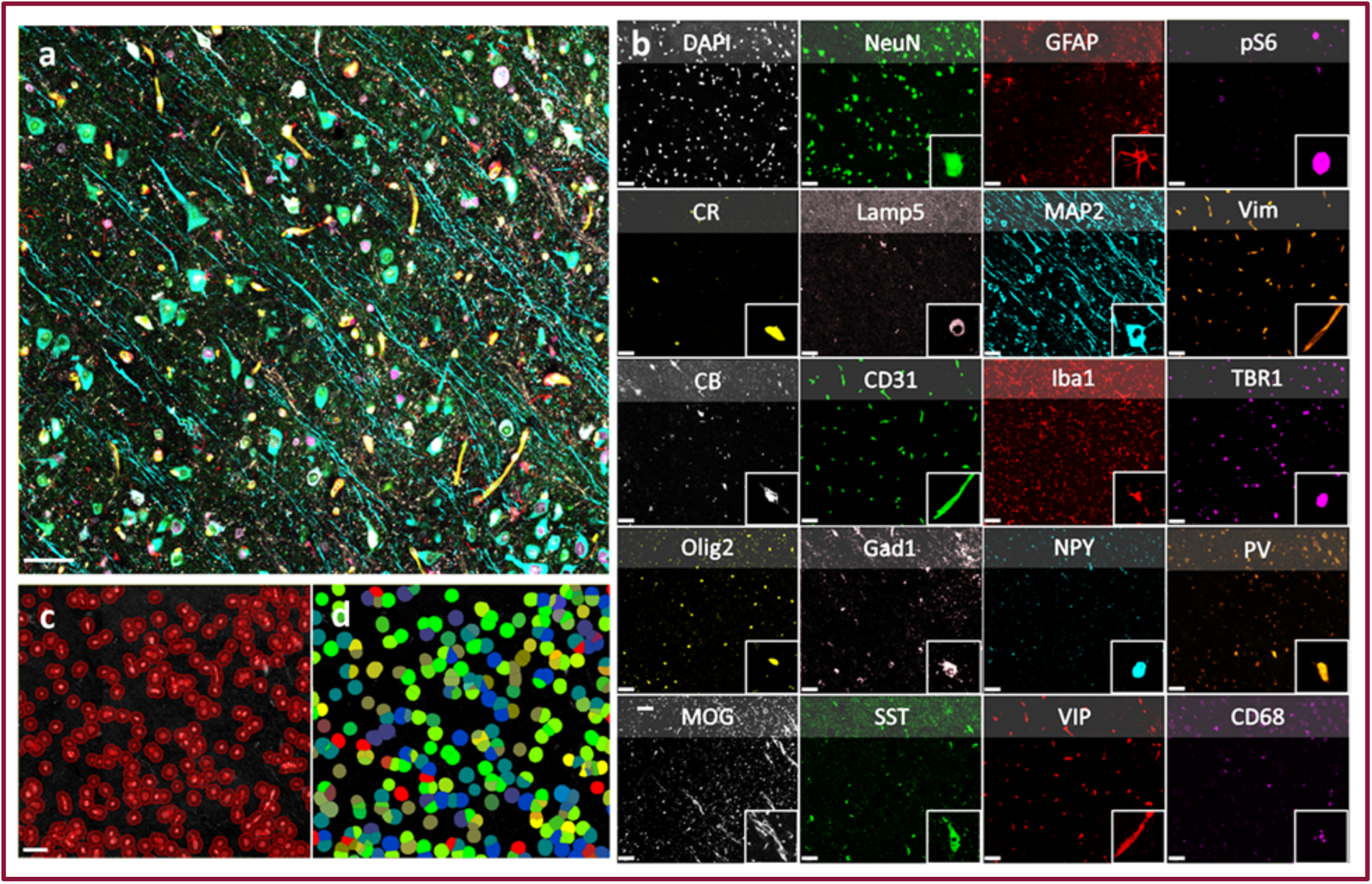
Novel multiplex neurobiology panel for characterising cell types in TSC. **a** Merge of the 20 channels. **b** Individual markers, with a zoom-in on each of the cell types. **c** Cell segmentation using StarDist. **d** Cell classification. Scale bar = 50 µm.

After the staining and image acquisition using the PhenoCycler-Fusion system, the images were first analysed using the QuPath platform ^21^. Cell segmentation was performed using a pretrained StarDist model, which is a dedicated nucleus segmentation algorithm^22^. StarDist represents each nucleus as a star convex polygon whose radial distances along fixed rays are predicted by a neural network, enabling accurate instance segmentation in crowded images^22,23^. This model detects nuclei and estimates cell boundaries by expanding from the nuclear position, allowing analyses of markers located in the cell body. We analysed 9 images from 7 different patients with a total of over 365 000 segmented cells (Fig. 2c). We randomly selected 27 regions representing the different areas of the grey matter, each of 250 000 μm2 of area to create a training image. Then, in conjunction with a neuropathologist, thresholds of positivity were determined for the markers, which were then merged into a composite classifier (Fig. 2d). This validated panel of multiplex cortical markers allowed us the first comprehensive single-cell resolution view of the TSC tissue architecture, enabling the systematic identification and analysis of cortical cell types and abnormal cells.

### Identity of TSC cell types

Recent studies have reported variable results as to the composition of tuberous tissue ^15,24^. Using our validated multiplex panel, our first goal was the characterisation of the various cell types present in the TSC tissue to help unravel the basis of epileptogenicity in this disease. Regarding the major classes of cortical cells, we used combinations of well-established markers to classify them as follows: neurons were defined by the presence of NeuN or MAP2; astrocytes by GFAP or Vim; microglia by Iba1; and oligodendrocytes by Olig2 (Fig. 3a). Quantification across segmented cells revealed a statistically significant increase in astrocytes, with TSC samples presenting an average of 29.10 ± 10.87% astrocytes compared to 5.43 ± 4.97% in controls (p = 0.0476). Microglia cells showed a mild, non-significant increase while oligodendrocytes and neurons showed a mild, non-significant decrease in TSC tissue compared to control cortex (Fig. 3b).

**Fig. 3.**
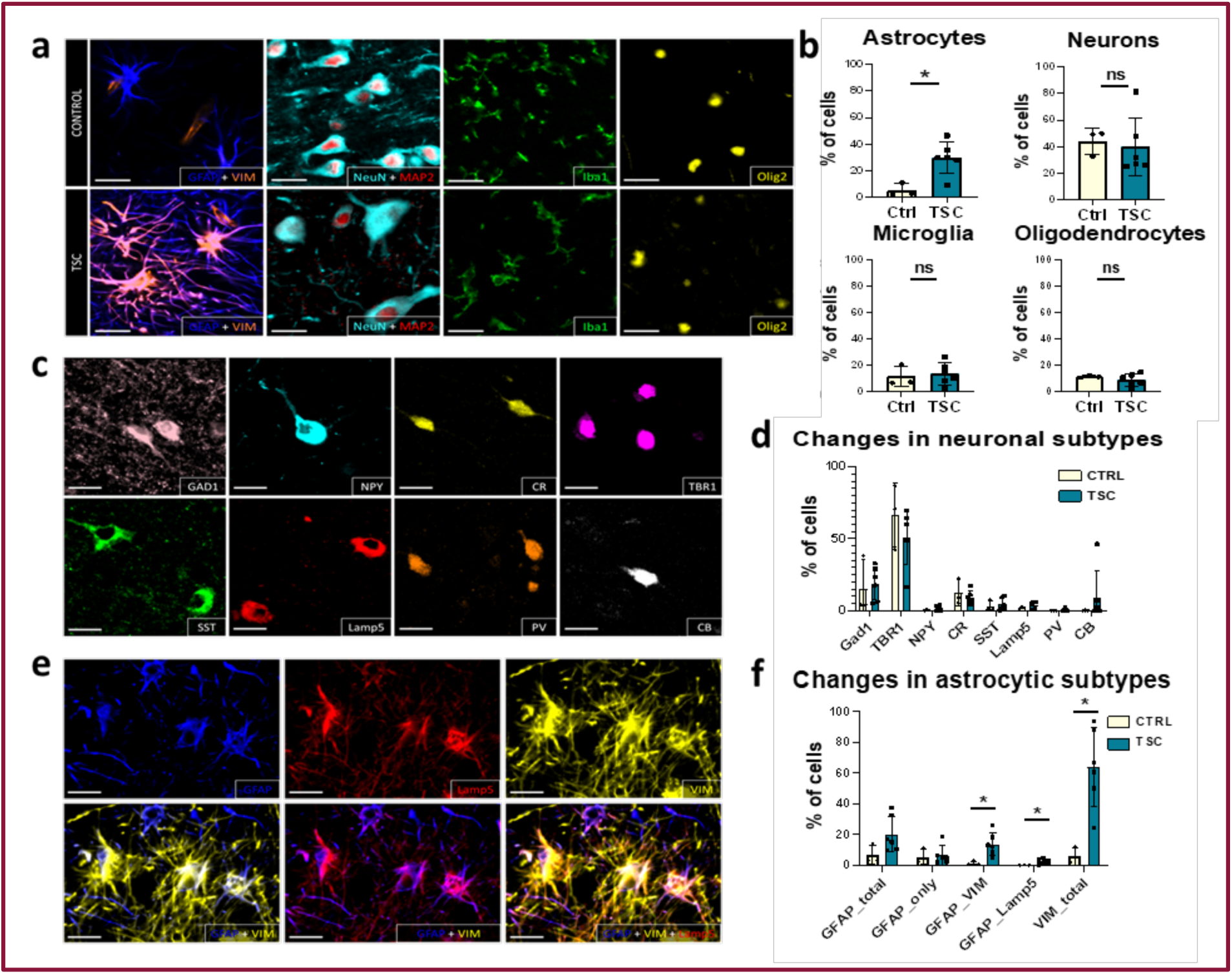
Quantification of cell types in TSC vs control. **a** Images from the general cell types found on the brain cortex of TSC and control patients: astrocytes Vim-positive or GFAP-positive, microglia Iba1-positive, neurons MAP2-positive or NeuN-positive and oligodendrocytes Olig2-positive. **B** Quantification of cell types. **c** Neuronal subtypes and **d** Quantification of neuronal cell types. **E** Astroglial subtypes in TSC vs control patients and **f** Quantification of astroglial subtypes. Scale bar = 50 µm. Statistical significance: ns = not significant; *p < 0.05.

To further resolve changes in neuronal composition, we analysed neuronal subtypes based on the expression of inhibitory neuron identity markers including CB, PV, SST, CR, NPY, Lamp5 and Gad1, as well as the excitatory neuron marker TBR1 (Fig. 3c). No statistically significant differences were observed between TSC and control samples for neuronal subtypes (Fig. 3d).

Next, we quantified astrocyte populations identified by GFAP, vimentin, and Lamp5 expression (Fig. 3e, f). Astrocytes stained only by vimentin were significantly increased in TSC tissue (63.78 ± 24.87%) compared to controls (5.58 ± 5.38%; p = 0.0238), suggesting an enrichment of immature or reactive glial populations. In contrast, more mature GFAP-positive astrocyte populations remained relatively stable, with a non-significant increase in GFAP-total astrocytes. However, GFAP/Vimentin double-positive astrocytes showed a significant increase from 1.32 ± 1.42% in controls to 12.43 ± 8.96% in TSC (p = 0.0238). There was also a modest but significant increase in GFAP/Lamp5-positive cells, from 0.14 ± 0.17% to 2.30 ± 1.53% (p = 0.0476). These results indicate a shift toward immature, metabolically active astrocyte phenotypes in the TSC cortex.

TSC is typically characterised by abnormally enlarged, dysplastic cells: BCs and DNs, that have been previously characterised as glial and neuronal-types, respectively ^18,25^. Both cell types can be identified based on their distinctly strong expression of the mTOR activation marker, pS6. The number of pS6-positive cells varies between patients, being in some cases scarce, usually restricted to the bottom of the sulcus, and throughout the white matter; while in other cases, they appear evenly distributed throughout the entire span of the cortical layers; its presence correlated with severity of the symptoms. Using the multiplex panel, DNs were defined as cells expressing pS6, NeuN or MAP2, and were further investigated for neuronal subtype lineage. While we found that their phenotype was primarily excitatory (35.54 ± 33.48% were TBR1-positive), a considerable proportion expressed the marker Gad1, indicating their inhibitory neuronal phenotype (15.10 ± 13.76%) (Fig. 4a). BCs on the other hand were defined as cells that were positive for pS6 expression and negative for NeuN or MAP2. We found that the vast majority of BCs express vimentin (95.90 ± 4.22%), as previously described ^18^, and 27.17 ± 25.60% were also positive for GFAP expression, and 14.05 ± 15.31% expressed Lamp5 (Fig. 4b), placing them in the glial phenotype with a strong tendency towards activated, immature subtypes.

**Fig. 4.**
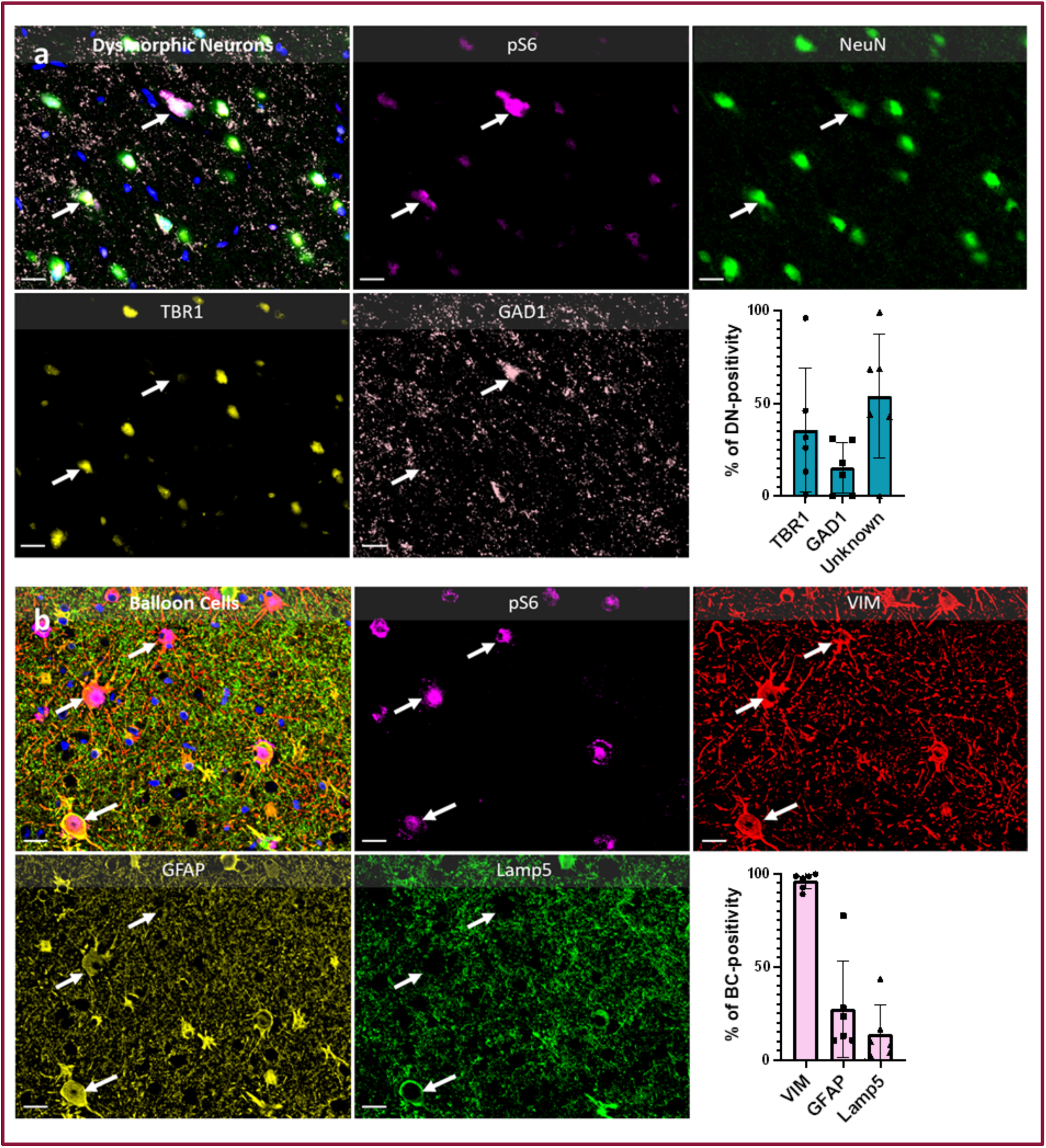
Abnormal cells found in TSC: BCs and DNs. **a** Marker positivity for DNs, expressing pS6, NeuN, TBR1 and GAD1; and quantification. **b** Marker positivity of BCs expressing pS6, Vim, GFAP and Lamp5. Arrows indicate the different combinations of markers found. Scale bar = 20 µm.

### Microtuber recognition and cell composition

Next, we investigated the cortical architecture disorganisation which has been associated with TSC cortical tissue. Microtubers are histopathologically abnormal structures first described as small, focal lesions in the cortex consisting of clusters of large abnormal cells and astrocytes (Fig. 5a) ^25^. Previous studies have demonstrated that these structures exhibit strong pS6 expression, reflecting hyperactivation of the mTOR signalling pathway ^26,27^. Additionally, Sosunov et al. provided a characterisation of reactive astrocytes within microtubers, differentiating them from those in larger tubers. Notably, their findings suggest that microtubers may represent the elementary unit of cortical tubers, accumulating over time and extending into perituberal areas ^16^. Thus, the cellular composition of these incipient microtubers could provide an insight into the mechanisms of formation of the large tuberous plaques that constitute TSC cortical lesions.

**Fig. 5.**
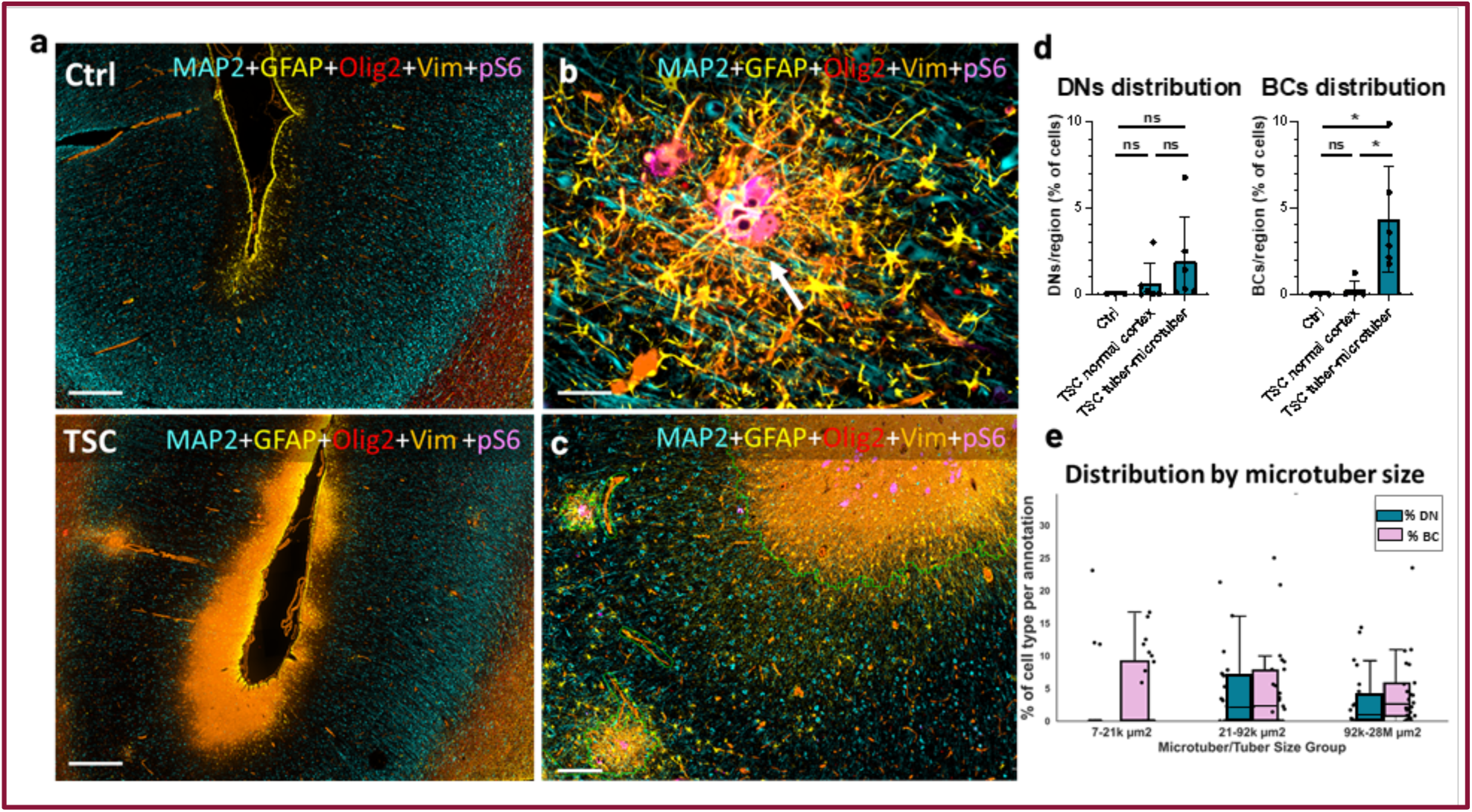
Microtuber formation in TSC tissue. **a** Widefield view of control case (upper panel) and a TSC case with microtuber/tuber-like areas shown by vimentin and pS6. **b** High magnification of a microtuber with pS6 (arrow) in the middle. **c** Microtuber recognition by the pixel classifier. **d** Quantification of abnormal cells inside and outside microtuber/tuber-like areas in control and TSC cases. **e** Distribution of DNs and BCs inside the microtubers segregated by groups of size by tertiles. Scale bars for a 500 µm, for b 50 µm; and for c 200 µm. Statistical significance: ns = not significant; *p < 0.05.

Here, we visually identified vimentin as a reliable marker for delineating microtuber boundaries, offering a precise means of recognising these pathological structures (Fig. 5b). To automate the microtuber recognition process, we trained a machine learning-based pixel classifier to detect vimentin-positive areas in the cortical grey matter, using training images from diverse regions across all patient sections to enhance robustness (Fig. 5c).

Upon validating the classifier, we applied it to all samples, successfully identifying microtubers and tuber-like structures in every TSC case, while none were detected in non-dysplastic, non-mTOR epileptic control cortex, confirming the specificity of our approach. We then investigated the presence and distribution of BCs and DNs in the microtuber/tuber-like areas and in the normal-looking cortex (Fig. 5). We found a significant enrichment of BCs within vimentin-positive tuberous formations (Fig. 5d), compared to both unaffected TSC cortex (p = 0.0107) and non-dysplastic controls (p = 0.0249), with BCs exclusively located within microtubers and none detected in regions lacking such glial architecture. Conversely, no significant difference was determined in the distribution of DNs between tuberous areas and normal-appearing cortex of TSC patients. To further investigate their spatial distribution of BCs and DNs in the incipient microtubers, we stratified vimentin-positive tuberous structures into size-based tertiles. This analysis revealed that BCs are the exclusive pS6-positive cell type present in the smallest microtubers, whereas both BCs and DNs are present and equally distributed in medium and larger structures (Fig. 5e). These findings support the hypothesis that BCs, rather than DNs, occupy the central niche of the immature gliosis and may play a primary role in driving its formation.

### Spatial analyses

To further explore the spatial organisation of pathological structures in TSC, we performed distance-based analyses to quantify the distribution of key cell types relative to microtuber/tuber-like areas. Using QuPath, we extracted distance-to-annotation values for each segmented cell, allowing us to measure the spatial relationships between microtubers and surrounding cellular populations. Using the microtuber annotation generated by the pixel classifier (Fig. 6a), we generated two heatmaps: one reflecting the distance to microtubers (Fig. 6b) and another showing distance to pS6 expression (Fig. 6c). While these heatmaps exhibited similar patterns, a difference was observed due to the presence of DNs expressing pS6 outside the microtuber boundaries.

**Fig. 6.**
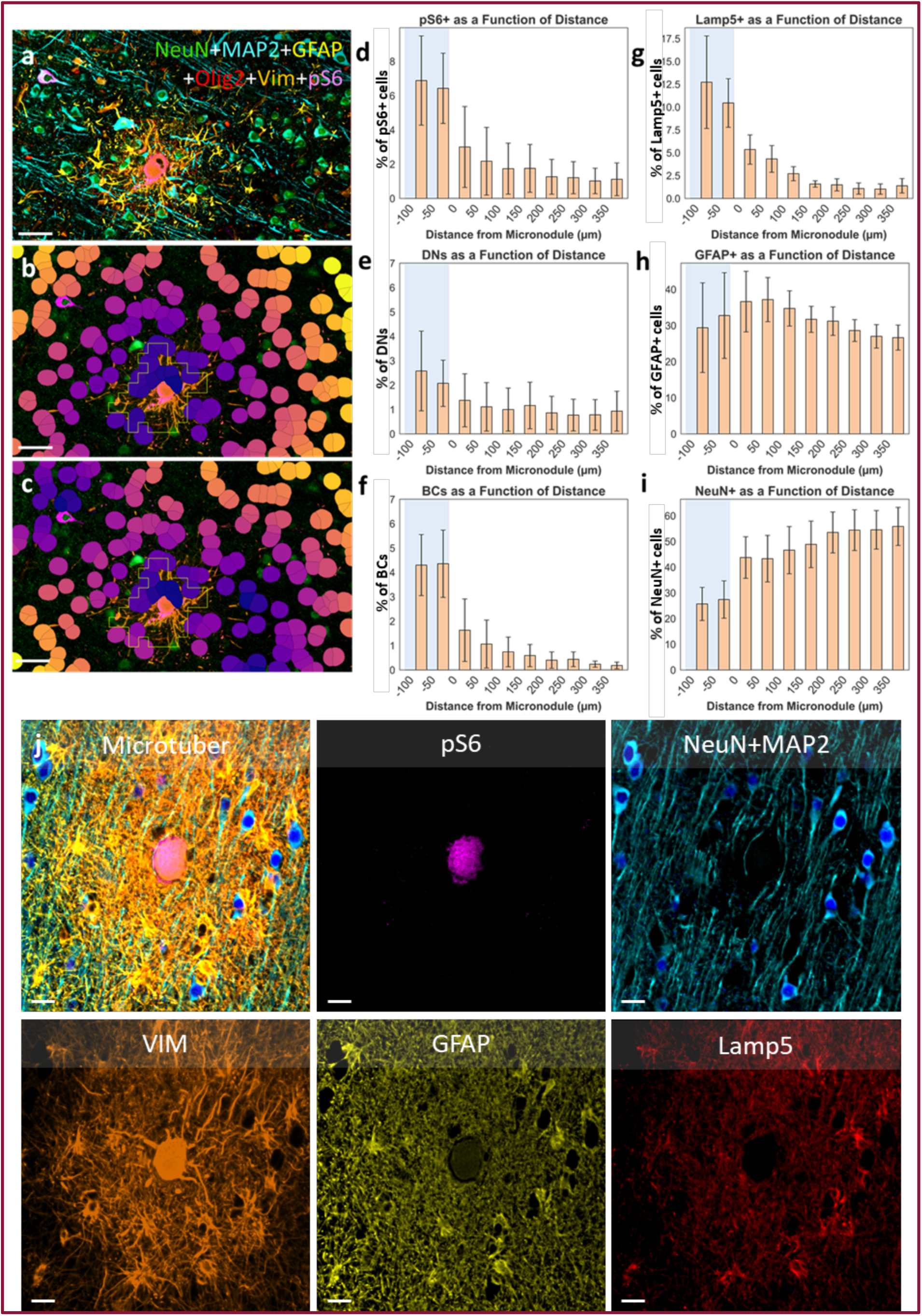
Spatial analyses of microtubers. **a** High magnification of a microtuber recognised by the pixel classifier. Scalebar = 50µm. **b** Heatmap of distance to microtubers. **c** Heatmap distance to pS6. **d** Quantification of pS6-positive cells in relation to the distance to microtubers. **e** Quantification of BCs in relation to the distance to microtubers. **f** Quantification of DNs in relation to the distance to microtubers. **g** Quantification of Lamp5-positive cells in relation to the distance to microtubers. **h** Quantification of GFAP-positive cells in relation to the distance to microtubers. **i** Quantification of NeuN-positive cells in relation to the distance to microtubers. **j** Microtuber characterisation. Scale bar = 20 µm.

We next quantified the proportion of specific markers as a function of distance from microtubers/tuber-like areas, using incremental bins of 25μm extending from -100μm (inside the microtuber) to 500μm outward (Fig. 6d-i). To support these spatial data with morphological evidence, we included a high-magnification multiplex image illustrating a microtuber archetype, where a central pS6- and vimentin-positive BC is surrounded by vimentin-positive glia and bordered by GFAP- and Lamp5-positive astrocytes, while NeuN- and MAP2-positive neurons are excluded from the core (Fig. 6j).

A progressive decline in the proportion of pS6-positive cells (BCs and DNs) was observed with increasing distance from the microtuber/tuber-like areas, although the decrease was not steep (Fig. 6d). This reflects the contribution of scattered pS6-positive DNs located outside the microtuber structures. When analysed separately, DNs showed a more gradual decline with distance (Fig. 6e). In contrast, BCs exhibited a sharp and localised distribution, with their density dropping rapidly beyond the microtuber boundary (Fig. 6f). These patterns indicate that BCs are tightly confined to microtuber cores, while DNs extend into the surrounding cortex. This spatial distinction is further exemplified in the representative image, where a central BC lies at the heart of the lesion, consistent with its restricted distribution (Fig. 6j).

To investigate the involvement of glial cells in microtuber pathology, we then quantified the different astrocytic populations. Lamp5-positive astrocytes were predominantly localised within and immediately adjacent to microtuber/tuber-like areas, with a distribution that closely mirrored that of pS6-positive cells (Fig. 6g). Interestingly, in several cases, Lamp5 expression was concentrated at the outer rim of the lesion, suggesting a defined boundary of glial remodelling (Fig. 6j). Meanwhile, GFAP-positive astrocytes displayed a unique pattern: their presence peaked in the microtuber/tuber-like peripheral zone but was comparable within or outside them (Fig. 6h, j). Together, these findings point to a stratified organization of reactive astrocytes within the microtuber environment.

Finally, to assess the impact of microtubers/tuber-like areas on cortical neuronal organisation, we quantified the distribution of NeuN-positive neurons relative to microtuber/tuber-like areas distance (Fig. 6i). In contrast to other markers, NeuN-positive cells density was all lowest inside microtubers/tuber-like areas, with a pronounced increase in density further from these structures. This supports the hypothesis that microtubers/tuber-like areas disrupt normal cortical architecture by displacing neurons or reducing their presence in affected areas (Fig. 6j).

### Unsupervised clustering reveals distinct cellular populations in TSC

Eventually, we performed unsupervised clustering on all segmented cells to augment the resolution of cellular subtype quantification. We identified a total of 38 cell clusters using binary marker expression features and cells were embedded in a low-dimensional space using UMAP. This approach revealed twelve distinct cellular clusters that were mainly found in all the samples (Fig. 7a, b). The clusters corresponded well to known cell identities, including excitatory neurons, inhibitory neurons, oligodendrocytes, astrocytes, microglia, and blood vessels, but also included pathologically relevant populations such as BCs, reactive astrocytes and DNs. Each cluster was annotated based on expression of known marker combinations, for instance, NeuN and MAP2 were highly expressed in excitatory and inhibitory neuronal clusters, while astrocytic populations were defined by GFAP and vimentin expression (Fig. 7c, d).

**Fig. 7.**
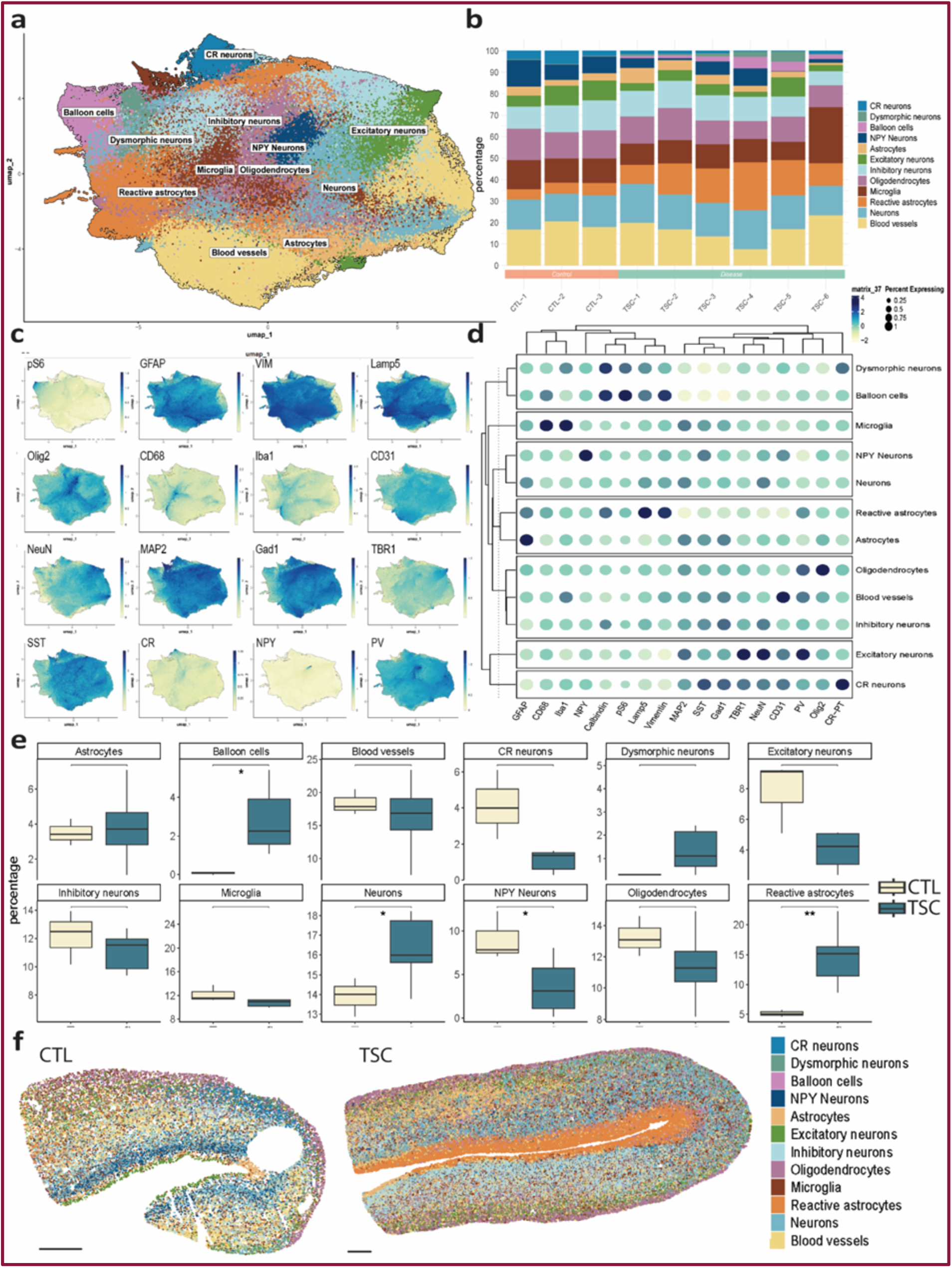
Unsupervised clustering of cell populations in TSC cortex. **a** UMAP of all segmented cells coloured by cluster identity revealing twelve major cell clusters. **b** Representation and percentages of the clusters on all samples. **c** Heatmap of marker expression on the UMAP. **d** Dot plot of marker expression across clusters. **e** Proportion of annotated clusters across TSC and control samples showed a significant increase of reactive astrocytes, BCs and not defined neurons and a reduction of NPY-positive neurons on TSC. **f** Spatial mapping of clusters onto representative tissue sections from TSC and control cases. Scale bar = 1 mm. Statistical significance: *p < 0.05; **p < 0.01.

We next compared the proportions of each cluster across TSC and control samples (Fig. 7e). While canonical cortical astrocytes (GFAP-positive) did not show a marked increase in TSC tissue, unsupervised clustering revealed a significant enrichment of reactive astrocyte populations, characterised by vimentin and/or Lamp5 expression. In contrast, oligodendrocyte and microglial proportions remained relatively stable between groups, consistent with the results of the supervised classification. Interestingly, we identified a neuronal cluster lacking clear excitatory or inhibitory marker expression, which appeared increased in TSC. However, given that both excitatory and inhibitory neuronal clusters were decreased, we interpret this as likely reflecting classification ambiguity rather than a biologically meaningful increase. Notably, NPY-positive neurons were significantly reduced in TSC samples, a finding that contrasts with the supervised analysis, highlighting how methodological differences can influence subtype detection. Finally, BCs and DNs were absent from control tissue but consistently identified in TSC samples, reinforcing their specificity to the disease context. BCs in particular were significantly enriched, mirroring the pattern observed in the supervised analysis.

Finally, using cell coordinates we mapped the cell clusters back onto the spatial tissue maps to examine their anatomical distribution (Fig. 7f, Supplementary Fig. 1). This revealed differences between control and TSC tissues with abnormal organisation where reactive astrocytes and BCs accumulated, corresponding to microtuber/tuber-like areas. Depending on the level of organisation, clusters showed disturbed spatial coherence in TSC compared to controls, indicating that the unsupervised clustering captured biologically relevant organisation.

### Niche analysis reveals a microtuber-associated cellular ecosystem

To further explore whether unsupervised clustering could delineate pathologically relevant microenvironments, we performed a niche-level analysis of cell populations. Niche analysis refers to the spatial profiling of cellular ecosystems within tissue, where cells are grouped into ‘niches’ based on their local neighbourhood composition and interactions rather than individual identity alone. By integrating cell type abundance, spatial proximity, and interaction networks, this approach enables the identification of recurring microenvironments that may underlie specific pathological features. This approach identified four distinct spatial niches across the TSC samples (Fig. 8a). Among these, niche 2 was enriched for reactive astrocytes and encompassed all balloon cells, while being spatially restricted to microtuber/tuber-like areas (Fig. 8b, c), thereby recapitulating the histopathological features of these structures. Network analyses showed reduced cell-cell interactions within niche 2 in several patients (Fig. 8d). In parallel, comparative quantification revealed that niche 2 displayed an increased mean distance to the nearest NeuN-positive neighbour across multiple clusters (Fig. 8e), further supporting the notion that neuronal disconnection is a defining feature of this environment.

**Fig. 8.**
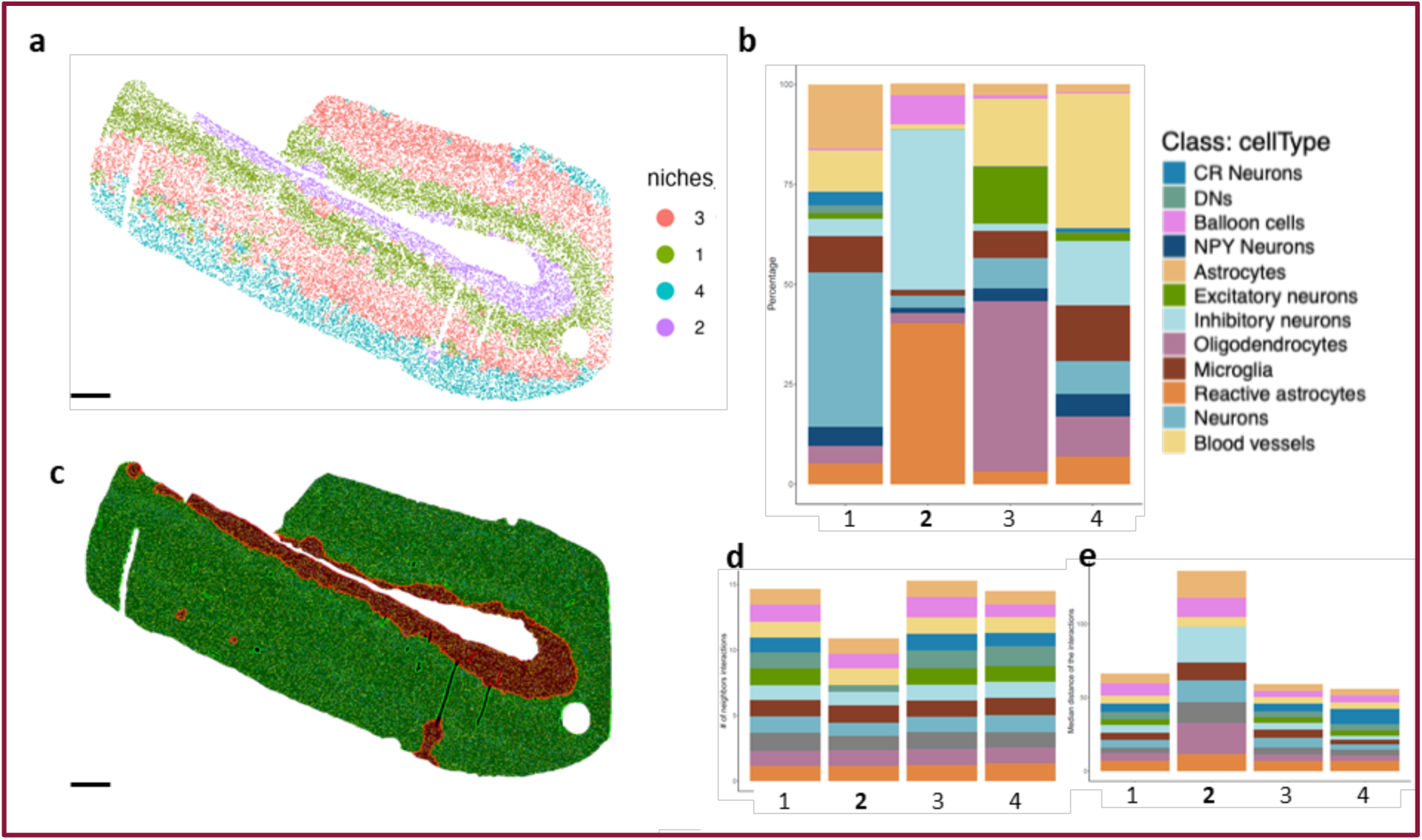
Niche analysis of TSC cortex. **a** Spatial mapping of four unsupervised niches across a representative TSC sample. **b** Proportional composition of cell types within each niche, showing an enrichment of reactive astrocytes and balloon cells in niche 2. **c** Automated recognition of microtuber/tuber-like areas (in red) by the QuPath pixel classifier, illustrating their spatial localisation within the sample. **d** Network analysis of cell-cell interactions across niches, revealing a reduction of interactions in niche 2 in some patients. **e** Quantification of mean distance to the nearest NeuN-positive cell across clusters, showing increased distances in niche 2. Scale bar = 800 µm.

Together, these findings suggest that niche 2 captures the pathological ecosystem of microtubers, in which BCs and reactive gliosis form a glial-dominated architecture that displaces neurons and disrupts local connectivity. This aligns with the clinical observation that seizure severity is linked to tuber/microtuber burden rather than a global imbalance of inhibitory and excitatory neurons, implicating reactive gliosis around balloon cells as a potential driver of epileptogenesis in TSC ^12,17^.

## Discussion

In this study we used for the first time in TSC multiplex spatial imaging combined with machine learning. Our main finding in cortical tubers is the identification of microtuber architecture, centred around balloon cells, which aggregate reactive immature glia and disrupt cortical organization. Above all, we provide a proof of concept that this method could be generalised and automated in neuropathology to decipher complex tissue architecture, enabling precise, high-throughput analysis of cell types and spatial patterns.

First, we developed a 19-marker panel optimised for cortical neurobiology. While pairing it with deep learning-based cell segmentation and pixel classification, we successfully classified over 365 000 individual cells, defined lesion boundaries based on vimentin expression, and explored spatial gradients of key cell types such as BCs, DNs, and glial subtypes.

Our method enabled an unprecedented comprehensive quantification and profiling of broad cortical cell classes. Compared to control tissue, TSC samples exhibited a significant increase in astrocyte populations, with slight but non-significant changes in oligodendrocytes, neurons and microglia. These population-level shifts reinforce prior findings that TSC pathology involves gliosis and glial remodelling ^17^. Further subclassification of astrocytes revealed their heterogeneity: Vimentin-positive astrocytes were significantly enriched in TSC tissue, suggesting an increase in immature or reactive glial states ^16^. In contrast, GFAP-positive mature astrocytes were relatively stable. This aligns with known mTOR-driven reactive gliosis and highlights the utility of multiplex markers in capturing glial heterogeneity that standard GFAP staining would miss.

Among neuronal subtypes, we observed no significant changes in the inhibitory or excitatory populations analysed, however we could reveal neuronal heterogeneity at a large scale. As an example, DNs in TSC are classically characterised as excitatory ^15^. However, our data reveals that a subset of DNs express also GAD1, in line with findings from FCD studies, where it was found that some DNs express inhibitory markers such as GAD67, calbindin or calretinin ^28^. These observations raise the possibility that DNs may arise from both excitatory and inhibitory lineages, or acquire mixed phenotypic features due to disrupted differentiation or fate specification under mTOR dysregulation.

Second, a key methodological achievement of our study was the development of vimentin-based pixel classification to reliably identify microtubers/tuber-like areas across patient sections and analyse cortical microarchitecture. This approach reduced the subjectivity of manual annotation and allowed for reproducible lesion detection in tissue with complex and heterogeneous architecture. The classifier showed high specificity, detecting microtuber-like structures in all TSC cases but none in controls.

This automated pipeline quantified the relative distribution of BCs and DNs inside and outside the identified lesions. BCs were significantly enriched within microtubers, whereas DNs showed a more diffuse distribution, appearing in tuber areas and extending beyond the lesion boundaries. This spatial pattern, reproducible across patients, supports the hypothesis that BCs are a central feature of microtuber pathology and may play a role in shaping their local environment. The spatial gradient of BCs, with a sharp decline outside microtuber boundaries, suggests their active disorganisational role. In contrast, the gradual decrease of DNs indicates their broader integration across pathological and adjacent tissue. These patterns imply distinct functional roles and origins for these two cell types within the TSC cortex, consistent with previous reports that link BCs to disrupted radial glial development, persistent glial activation and local immune activation. As in FCD type IIb, BC load correlates with complement activation, antigen presentation, and T-cell infiltration, implicating these cells in local inflammation^29^. Although our panel lacked immune markers, the presence of pS6-positive BC cores encircled by reactive vimentin-positive glia likely represents the structural correlate of such inflammatory niches in TSC.

Eventually, we developed an automated unsupervised method that could segregate cell subtypes in distinct population clusters and interpolate their spatial distribution. Cell quantification was comparable with the supervised approach, while reactive astrocytes aggregates were detected by the algorithm and corresponded to the micro tuber areas.

In addition, our unsupervised niche analysis independently confirmed that microtuber/tuber-like structures constitute a distinct cellular ecosystem. Niche 2 was uniquely defined by the presence of BCs and reactive astrocytes, while being spatially restricted to microtuber regions identified by the QuPath classifier. This niche displayed both an increased mean distance to the nearest NeuN-positive neighbour and reduced neuronal density, pointing to neuronal displacement and local network disruption. These findings are consistent with clinical observations that seizure severity in TSC correlates with tuber and microtuber burden rather than a global imbalance of excitatory and inhibitory neurons ^12,16^. They also reinforce the concept that reactive gliosis centred around BCs is a key participant in epileptogenesis^17,26^. Understanding the mechanisms that drive the formation of this gliotic niche may therefore open avenues for therapeutic strategies targeting glial-neuronal interactions in TSC.

Taken together, multiplex imaging combined with machine learning can be widely generalised to the neuropathology of MCDs, providing insight into both cell quantification and spatial architecture. By resolving a microtuber centred ecosystem driven by BCs and reactive astrocytes and quantifying its neuronal depletion, we provide a mechanistic scaffold for tuber formation in TSC. The multiplex imaging and machine learning workflow that enabled this result scales to large cohorts and other cortical malformations, establishing a practical path to routine, quantitative neuropathology.

## Materials and methods

### Patient and tissue

A comprehensive presurgical evaluation was performed with EEG-video monitoring, 3T MRI, neuropsychological evaluation, genetic testing, and when required 18FDG-PET or intracranial EEG recordings with depth electrodes (SEEG). The surgery was not modified for the study and consisted in “en bloc resections” to avoid excessive manipulations of the tissue as previously reported ^30^.

Control cortex was obtained as part of temporal lobe resections where both the temporal pole and mesial structures needed to be resected. This cortex proved to be spared of pathological process during histological analysis.

Patients’ parents or legal guardian gave informed written consent for the storage, and research on biological specimens provided as part of treatment. The biological collection stored in IHU Imagine / Necker Hospital and the study are declared respectively under n° D2009-955, and n° ANR 20-CE17-003.

### Histopathological diagnosis

The tissues have been selected after a standard histopathological diagnosis, carried out according to the state of the art by a neuropathologist. The regions of interest were selected by the same neuropathologist. Standard immunohistochemistry confirmed the presence or absence of BCs, DNs, subpial gliosis and GFAP positive micronodules.

Histopathological diagnosis was performed on 3-μm-thick slides stained with Hematoxylin-Phloxine-Saffron (HPS) according standard protocol on an automated stainer (HistoCore Spectra, Leica) and 3-μm-thick slides immunostained using an automated immunostainer (Dako Omnis, Glostrup, Denmark) with the following antibodies: Microtubule Associated Protein (MAP-2; Sigma ; HM-2 ; 1/20 000), Neuronal Nuclei (NeuN; Eurobio ; A 60 ; 1/1000), Glial Fibrillary Protein (GFAP; Dako ; 6F2 ; 1/200); Phospho-S6 (Cell signaling ; Polyclonal ; 1/400).

### Creation of the panel

To study the architecture of the cortical tissue and the identity of the cell types in the TSC pathology, an antibody panel was designed (Supplementary Table 2) following the knowledge on neurobiology markers in the literature.

### Conjugation of antibodies

All antibodies selected for the panel were purchased from commercial vendors and subsequently conjugated to custom oligo barcodes following the Akoya Biosciences’ conjugation kit protocol (Conjugation kit, #7000009; Akoya). Briefly, each antibody is linked to a unique oligonucleotide sequence after a reduction step that exposes the thiol groups, and once purified, they are kept at 4°C until use. This oligonucleotide sequence is called barcode, and it will be recognised by a complementary oligonucleotide sequence linked to a fluorophore ^31^.

### Multiplex imaging

#### Sample processing

The fixation and staining protocol were performed according to Akoya User Manual of the PhenoCycler-Fusion: after baking the slides overnight at 60°C, they undergo an automated deparaffination using xylene and sequentially decreasing concentrations of ethanol to rehydrate the tissue. The staining was performed using conjugated antibodies from Akoya Biosciences at different concentrations and custom-conjugated antibodies from other companies (Supplementary Table 2) following the protocol from Akoya Biosciences.

#### Reporter plate preparation and imaging

The PhenoCycler-Fusion assay was performed following Akoya Biosciences’ recommendations: each cycle consists of 4 fluorescent channels (3 for antibody visualisation and 1 for DAPI nuclear stain) and they were prepared in a 96 well/plate with 1 well per cycle containing the fluorescent oligonucleotides (reporters) and only buffer with DAPI staining for the blank cycles. These blank cycles are the first and the last one, and the system uses them for the focus and subtraction of autofluorescence in the background.

For imaging, the slides were assembled onto a custom-designed plate holder, named flow cell (#240204, Akoya Biosciences) and then imaged on the PhenoCycler-Fusion using the optimised exposure settings described in table 1. The DAPI nuclear stain (concentration 1:600) was imaged in each cycle at an exposure time of 10ms. The process consists in an automated and integrated process of fluidics that performs hybridisations of the reporters, imaging of the tissue and dehybridisation in each cycle. At the end, the system creates a single multichannel .qptiff image which then was imported into the QuPath software ^21^.

### Bioinformatics analyses

#### Image processing in QuPath

Image analysis followed the workflow described in a prior Imaging Mass Cytometry (IMC) study of kidney inflammation that used a comparable number of markers^5^. All images were imported into a QuPath project. To construct a robust pipeline, we assembled a composite training image by stitching together multiple 500 × 500 µm regions randomly sampled from grey matter across cases to capture the diversity of cortical structures. This composite was then used to optimize cell segmentation, define marker positivity thresholds, train the vimentin-based pixel classifier for microtuber recognition, and derive spatial features. All steps were performed under the supervision of a neuropathologist.

#### Cell segmentation

StarDist is a deep learning-based segmentation algorithm that uses a star-convex shape representation to achieve accurate and reliable nucleus segmentation in QuPath ^21,32^. We used the script *Multimodal StarDist Segmentation.groovy* with the slight modification that we added a line to define the maximum nuclear area to 210 µm^2^ and we set the minimum nuclear area to 10 µm^2^ to avoid detecting artifacts ^23^. The segmentation was based on the use of the DAPI channel using a 10 µm expansion, 0.6 detection probability threshold and 1st to 99th percentile normalisation and the pretrained model dsb2018_heavy_augment.pb.

#### Cell Classification

Under the supervision of a pathologist, the thresholds of positivity were set for 16 of the markers. For 9 of the markers, single measurement classifiers were set based on mean or standard deviation intensity in either the nucleus or the cell depending on each marker. While for the other 7, object classifiers were needed for the complexity or the markers and variability of the samples. Final thresholds and features used for each individual channel detailed in Supplementary Table 3. Finally, a composite classifier of all the markers was created and applied to the segmented cells.

### Pixel classifier for microtuber recognition

Vimentin was identified as a reliable marker for detecting microtuber or tuber-like areas in the grey matter based on histopathological evaluation. Another training image was created with different areas that contained a representative view of all the areas. A pixel classifier was trained using the machine learning model artificial neural networks (ANN) and the vimentin channel. Along with the following features: Gaussian, Laplacian of Gaussian, Weighted Deviation and Gradient Magnitude; and all the scales (0.5, 1, 2, 4, 8).

The pixel classifier created 2 types of annotation objects that were inserted into the image annotation hierarchy along with the segmented cells. From this moment, the cells contained information about their “Classification”, positivity for markers and their “Parent”, whether or not they were part of the microtuber.

#### Single-cell spatial features

Spatial data was generated for each of the cells: annotation distances, cell distances and Delaunay triangulation. Annotation distances is the distance of each cell to the nearest annotation. Cell distances is the distance of each cell to the nearest cell positive for each of the markers with a maximum distance cap of 200 µm. Delaunay triangulation is a neighbourhood-based analyses that identifies adjacent cells within a defined distance threshold and computes descriptive statistics for features of those neighbouring cells. In this study, Delaunay triangulation was computed using a 20 µm clustering threshold and restricted to neighbouring cells sharing the same classification.

### Script generation and data export

The workflow was consolidated into a single script and applied to all original images, focusing on grey matter regions defined histopathologically. The resulting data was exported into two CSV files, a per-cell dataset, where each row represents an individual cell and each column corresponds to a measured feature; and a per-annotation dataset, where each row corresponds to an annotation generated by the pixel classifier (Microtuber/Tuber-like area or normal cortex).

### Data processing and quantification of marker positivity

Building on the multiplexed analysis workflow of Alexander and Zaidi et al., the analysis of the exported datasets was performed using a set of Python scripts. These scripts were used to calculate the percentage of single- and double-positive cells corresponding to the major cortical cell types, including neuronal and astrocytic subtypes, and abnormal cells in both control and TSC samples. The same approach was applied to quantify marker expression within abnormal cell types including the percentages of positivity for each marker. Annotation-level measurements were also used to compare the distribution of abnormal cells in microtubers stratified by size (using tertile-based classification).

#### Spatial quantification

The signed distance from each marker or cell-of-interest to the nearest microtuber boundary was binned in 50 µm increments, ranging from -100 µm (inside) to +400 µm (outside). The percentage of marker-positive cells was calculated per sample and distance bin, to highlight the relative spatial position of cells around microtubers.

#### Statistical analyses

Statistical analyses were conducted using Python (version 3.10) and GraphPad Prism (version 8). For comparisons between TSC and control, normality was assessed using the Shapiro–Wilk test and variance equality using Levene’s test. When both assumptions were met, an unpaired two-tailed Student’s t-test was used. Otherwise, the two-sided Mann–Whitney U test was applied.

For datasets involving more than two groups, statistical comparisons were first assessed using the Kruskal–Wallis test. When a significant effect was detected, pairwise comparisons were conducted using the Mann–Whitney U test. For parametric data, one-way ANOVA followed by Tukey’s multiple comparisons test was applied using GraphPad Prism. Significance thresholds were defined as *p < 0.05, **p < 0.01, and ***p < 0.001.

Exact p-values were reported unless otherwise stated, and a p-value < 0.05 was considered statistically significant. The unit of analysis was the individual patient. In two cases, two anatomically distinct cortical regions were analysed per patient to increase statistical power; these were treated as independent observations in the group-level comparisons. Results are presented as mean ± standard deviation (SD), unless otherwise specified.

#### Unsupervised clustering

The csv files were loaded into Seurat (v5.10) using a modified version of seurat ReadAkoya function. The changes to the code were taken directly from the issues section of seurat git repository (https://github.com/satijalab/seurat/issues/7792).

Cells with low counts (nCount_Spatial < 5), low number of protein signal (nFeature_Spatial < 5) and a detection probability lower that 0.7 were removed. The data from the remaining cells were centered log ratio normalised using the CLR implementation and the effect of different number of counts among samples was removed during scaling. Batch effect between samples was corrected using Seurat’s FindIntegratedAnchors, and the samples were integrated using the harmony reduction. On the integrated dataset, dimensionality reduction was performed using principal component (PC) analysis computed on the first 20 PC. FindNeighbors was used to compute the k-nearest neighbor graph on the low-dimensional embedding, and FindClusters was used to cluster the cells with multiple resolutions (from 0.4 to 1.2). Finally, the reference resolution with r=1 was considered to better reflect the expected cell populations, and it was used for visualisation and all downstream analyses. UMAPprojection was calculated using Seurat RunUMAP function with the metrics argument set to Euclidean. Cell type labels were assigned based on the classification performed with QuPath. All clusters were annotated, and 329018 cells were kept.

## Code availability

Custom code used for data analysis in this study is available from the corresponding author upon request and at https://github.com/MarkZaidi/Kabashi-Lab.git

## Author contributions

B.A., B.C., M.v.V., and R.W. have made substantial contributions to the conception, design, financial acquisition, analysis, and interpretation of the data, drafting of the work, and revised the manuscript. R.G., T.K., E.B., W.N., and C.O. have made substantial contributions to the conception, design, analysis, and interpretation of the data, drafting of the work, and substantially revised the manuscript. F.B., M.v.W., B.S., P.C., J.P., E.A., and W.J.W. have made substantial contributions to the conception, design, analysis of the data, drafting, and revision of the manuscript. All authors have approved the submitted version and agree to be personally accountable for the author’s own contributions and to ensure that questions related to the accuracy or integrity of any part of the work, even ones in which the author was not personally involved, are appropriately investigated, resolved, and the resolution documented in the literature.

