## Supplementary data for "Multiplex imaging combined to machine learning enable automated profiling of cortical malformations: applications in tuberous sclerosis complex"

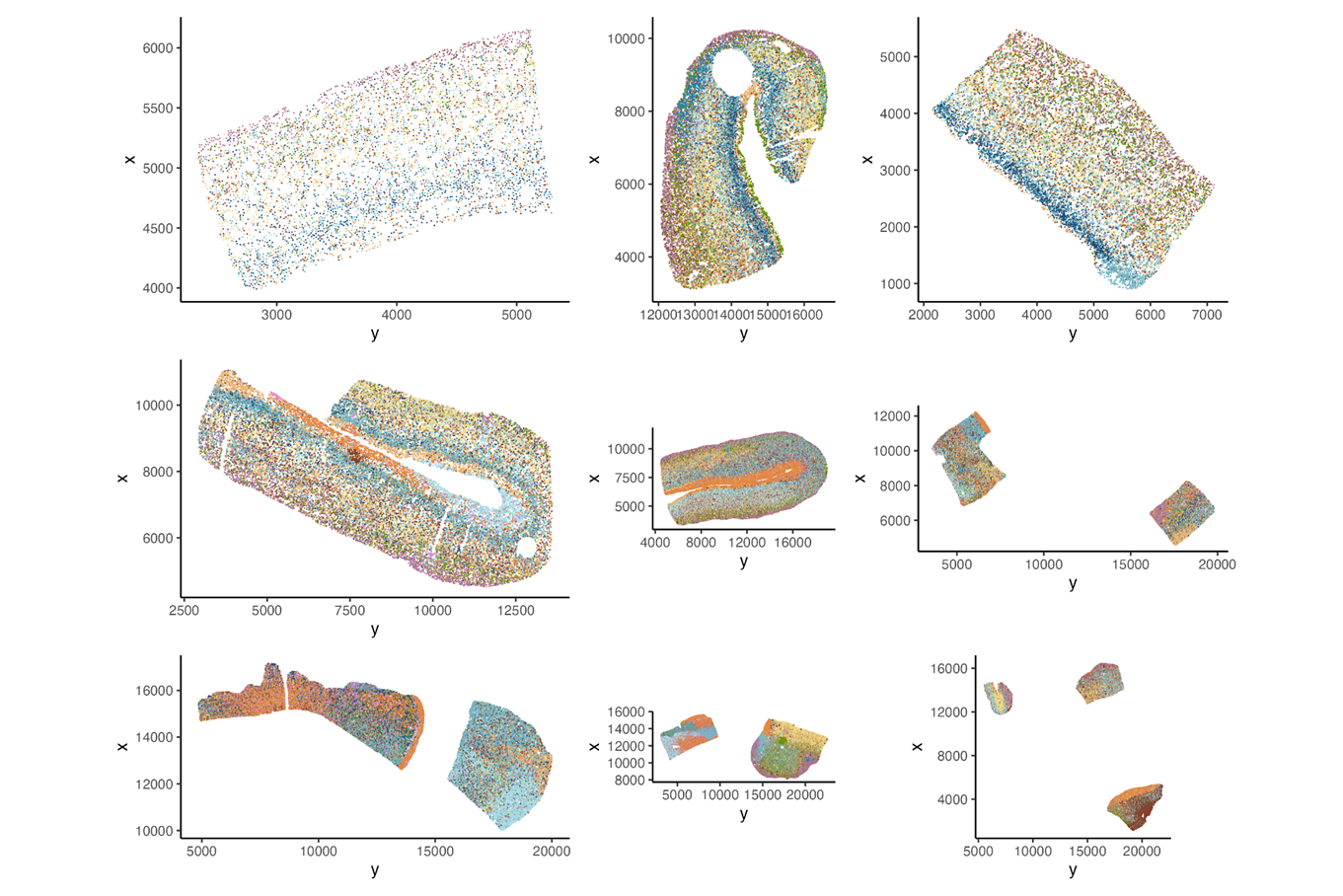

**Supplementary Fig. 1** Spatial mapping of clusters on the tissue of all samples

**Supplementary Table 1.** Patients’ data

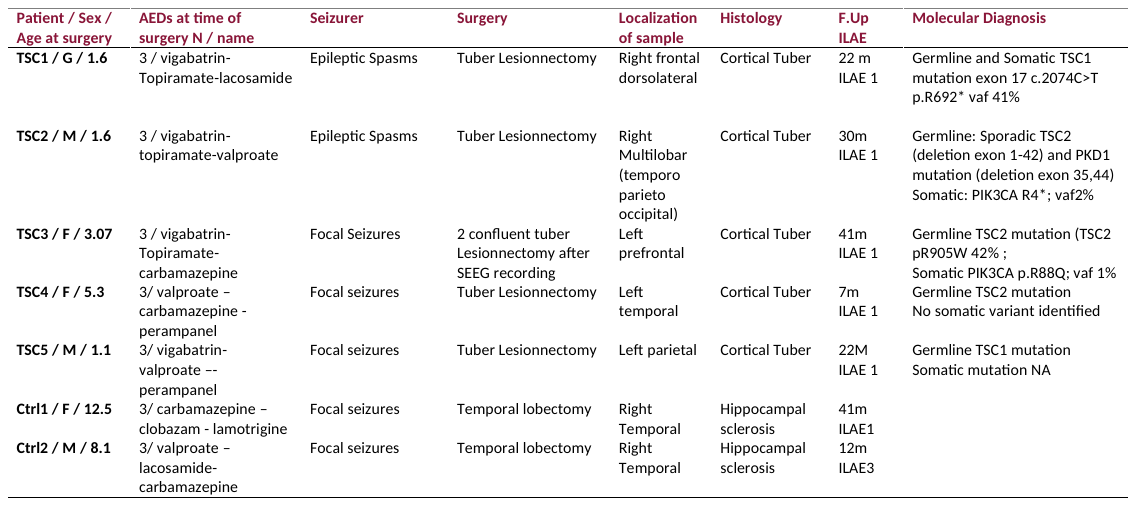

**Supplementary Table 2.** Antibody panel used for the PhenoCycler-Fusion

| **Antibody** | **Concentration** | **Exposure time (ms)** | **Commercial brand** | **Reference** |
| --- | --- | --- | --- | --- |
| CD31 | 1:200 | 150 | Akoya Biosciences | 4250104 |
| CD68 | 1:200 | 80 | Akoya Biosciences | 4550113 |
| Vimentin | 1:200 | 120 | Akoya Biosciences | 4450050 |
| NeuN | 1:200 | 200 | BioLegend | 834501 |
| GFAP | 1:200 | 80 | Clinisciences | AM1870-ev |
| Calbindin-D28k | 1:100 | 150 | Fisher Scientific | 15674295 |
| Lamp5 | 1:100 | 200 | Fisher Scientific | 15587676 |
| pS6 | 1:100 | 250 | Merck | 07-2113 |
| Calretinin | 1:50 | 180 | Proteintech | 66496-1-Ig |
| Gad1 | 1:100 | 80 | Proteintech | 67648-1-Ig |
| Iba1 | 1:200 | 120 | Proteintech | 10904-1-AP |
| MAP2 | 1:200 | 100 | Proteintech | 17490-1-AP |
| MOG | 1:200 | 150 | Proteintech | 12690-1-AP |
| NPY | 1:100 | 180 | Proteintech | 12833-1-AP |
| Olig2 | 1:200 | 150 | Proteintech | 13999-1-AP |
| Somatostatin | 1:100 | 200 | Proteintech | 17512-1-AP |
| TBR1 | 1:100 | 200 | Proteintech | 20932-1-AP |
| VIP | 1:200 | 180 | Proteintech | 16233-1-AP |
| Parvalbumin | 1:50 | 200 | Swant | 235-PUR |

**Supplementary Table 3.** Final thresholds and features of the algorithm used for each individual marker

| Antibody | Staining | Algorithm | Feature |
| --- | --- | --- | --- |
| CD31 | Cellular | Object classifier | All CD31 |
| Vim | Cellular + projections | Single classifier | Cell SD = 23,1245 |
| NeuN | Nuclear | Single classifier | Nuclear Mean = 15 |
| GFAP | Cellular + projections | Object classifier | All GFAP |
| CB | Cellular | Single classifier | Cell SD = 2,9 |
| Lamp5 | Cellular + projections | Object classifier | All Lamp5 |
| pS6 | Cellular | Single classifier | Cell SD = 3,4 |
| CR | Cellular | Single classifier | Nuclear Mean = 3 |
| Gad1 | Cellular or membranous | Object classifier | All GAD1 |
| Iba1 | Cellular + projections | Single classifier | Nuclear Mean = 6,6 |
| MAP2 | Cellular + projections | Object classifier | All NeuN + MAP2 |
| NPY | Cellular + projections | Single classifier | Nuclear Mean = 9 |
| Olig2 | Cellular | Single classifier | Nuclear SD = 7 |
| SST | Cellular | Object classifier | All SST |
| TBR1 | Nuclear | Single classifier | Nuclear SD |
| PV | Cellular | Object classifier | All PV + Olig2+Nucleus area, length, max, min |
